# Learning Clique Subgraphs in Structural Brain Network Classification with Application to Crystallized Cognition

**DOI:** 10.1101/2020.05.26.116475

**Authors:** Lu Wang, Feng Vankee Lin, Martin Cole, Zhengwu Zhang

**Affiliations:** Department of Statistics, Central South University; Elaine C. Hubbard Center for Nursing Research On Aging, School of Nursing, University of Rochester Medical Center; Department of Psychiatry, School of Medicine and Dentistry, University of Rochester Medical Center; Department of Brain and Cognitive Sciences, University of Rochester; Department of Neuroscience, University of Rochester Medical Center; Department of Neurology, School of Medicine and Dentistry, University of Rochester Medical Center; Department of Biostatistics and Computational Biology, University of Rochester Medical Center

**Keywords:** network classification, signal subgraph learning, clique subgraphs, structural brain networks, symmetric bilinear logistic regression

## Abstract

Structural brain networks constructed from diffusion MRI are important biomarkers for understanding human brain structure and its relation to cognitive functioning. There is increasing interest in learning differences in structural brain networks between groups of subjects in neuroimaging studies, leading to a variable selection problem in network classification. Traditional methods often use independent edgewise tests or unstructured generalized linear model (GLM) with regularization on vectorized networks to select edges distinguishing the groups, which ignore the network structure and make the results hard to interpret. In this paper, we develop a symmetric bilinear logistic regression (SBLR) with elastic-net penalty to identify a set of small clique subgraphs in network classification. Clique subgraphs, consisting of all the interconnections among a subset of brain regions, have appealing neurological interpretations as they may correspond to some anatomical circuits in the brain related to the outcome. We apply this method to study differences in the structural connectome between adolescents with high and low crystallized cognitive ability, using the crystallized cognition composite score, picture vocabulary and oral reading recognition tests from NIH Toolbox. A few clique subgraphs containing several small sets of brain regions are identified between different levels of functioning, indicating their importance in crystallized cognition.

## 1. Introduction

Recent advances in magnetic resonance imaging (MRI) techniques enable us to noninvasively probe the human brain at higher resolutions than ever before [Glasser et al., 2016] and reconstruct connectomes with distinct physiological meanings [Park and Friston, 2013]. Among them, the diffusion MRI (dMRI) infers the locations and directions of white matter (WM) fiber tracts via measuring the water molecular movement along major fiber bundles in WM. Diffusion MRI data are now collected in almost all major cohort-based neuroimaging studies, e.g., the Human Connectome Project [Van Essen et al., 2013], the UK Biobank [Miller et al., 2016] and the Adolescent Brain Cognitive Development (ABCD) Study [Casey et al., 2018]. Structural connectivity (SC) analysis is among the most important applications of dMRI [Park and Friston, 2013; Zhao et al., 2015; Yeh et al., 2016; Zhang et al., 2018], where individual-level microstructural brain networks are constructed to delineate anatomical connections between brain regions. Figure 1 illustrates the pipeline we used for extracting SC [Zhang et al., 2018] (Figure 1a) and an SC matrix extracted from one subject in the ABCD data (Figure 1b).

**Figure 1:**
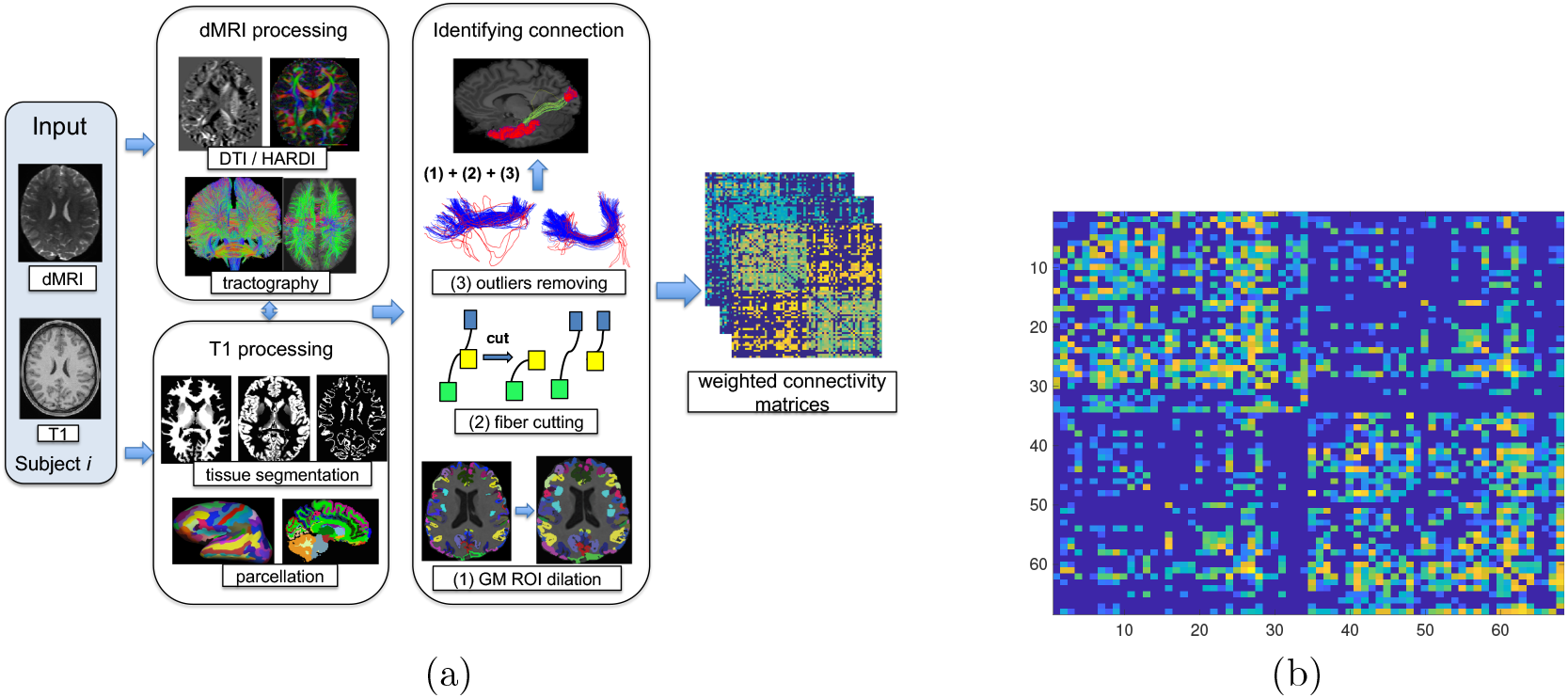
(a) shows the pipeline we used to extract the network and (b) shows a structural brain network extracted from the ABCD data.

Brain network classification and identification of predictive subnetworks are probably among the most important applications of SC into the mechanistic understanding of neuroscience phenomena. One typical approach to this network classification and variable selection problem is to treat all the connections of SC as a long feature vector, and apply existing classification methods for vectors, such as generalized linear models with *L*_1_ or elastic-net penalty [Zou and Hastie, 2005], support vector machines [Zhu et al., 2004] etc. Another popular approach in the neuroscience literature is to perform massive univariate tests at each edge with multiple testing correction to identify edges that are significantly different between two groups. While these methods may have good prediction, they ignore the network nature of the data and do not guarantee any structure among the selected individual edges, making the results hard to interpret.

Other feature extraction approaches often employ a two-step procedure, where some unsupervised dimension reduction is first applied on the networks, and then a regression model is fitted on the extracted low-dimensional features. Zhang et al. [2019] proposed to use tensor decomposition approach to map networks to low-dimensional vectors. Simpson et al. [2011] proposed to reduce the networks to some topological summary measures such as clustering coefficient, network density, etc. However, such connectome simplification leads to an enormous loss of information and brings troubles in truly understanding which part of the brain network is responsible for the group difference.

There are a few advanced statistical methods considering the network structure when analyzing group differences in connectome. Vogelstein et al. [2012] proposed to search for a minimum set of vertices and edges distinguishing groups, but this method only applies to binary edges and involves solving a combinatorial optimization problem. Durante et al. [2018] developed a method for testing global and local changes in SC between groups, which gains power by accounting for dependence structure among edges through a Bayesian nonparametric modeling of the networks. Relión et al. [2019] proposed a graph classification method that uses edge weights as predictors and incorporates *L*_1_ and group lasso penalty to penalize both the number of edges and the number of nodes selected.

We developed a symmetric bilinear logistic regression (SBLR) model with elastic-net penalty to select a set of small clique signal subgraphs in network classification. A clique subgraph in graph theory refers to a subset of nodes of an undirected graph such that every two different nodes in the clique are connected [Seidman and Foster, 1978]. SBLR puts symmetry constraint on the coefficient matrix in the logistic regression, because the adjacency matrices of structural brain networks are symmetric. The novelties and significance of our approach reside on:

- The small clique subgraphs identified by our method have appealing interpretations in neuroscience field. Clique subgraphs (containing all the interconnections among a subset of nodes) potentially aligns with the physiological meaning of subgraphs that should be strongly interrelated in order to provide the most efficient neural support of a behavior [Bassett and Sporns, 2017]. Despite the clique structure imposed on the selected subgraphs, our model maintains the flexibility of identifying individual edges in network classification and good classification rate.
- The elastic-net penalty penalizes the size of each identified clique subgraph, producing a parsimonious representation of differences in brain connectome. We develop a coordinate descent algorithm for model estimation where analytical solutions are derived for a sequence of conditional convex optimizations. The code for implementing the algorithm is publicly available at https://github.com/wangronglu/SBLR.
- The SBLR approach is applied to examine structural network classification for crystallized intelligence among 4213 right-handed adolescents aged 9-10 years in the ABCD study. Emerging literature suggests unique roles of white matter in supporting general cognition and differentiating crystallized and fluid intelligence [Simpson-Kent et al., 2020; Penke et al., 2012; Góngora et al., 2020]. Of note, age 9-10 represents a critical change point for the relationship between white matter and crystallized intelligence [Simpson-Kent et al., 2020]. Thus far, however, no study has specified the role of SC in crystallized intelligence of adolescents. We extensively analyzed SCs of subjects in ABCD with high vs. low crystallized intelligence using our SBLR model, aiming at learning interpretable SC-based brain connectome maps that can differentiate the levels of crystallized intelligence.

The rest of the paper is organized as follows. Section 2 introduces the data preprocessing steps, our method and the algorithm for model estimation. Section 3 presents simulation studies demonstrating good performance of SBLR in recovering true clique signal subgraphs compared with other methods. Application of this method to ABCD data in Section 4 shows coherent subgraphs of crystallized intelligence across composite score and individual domains. We conclude with a brief discussion in Section 5.

## 2. Methods

### 2.1. ABCD Data Preprocessing

We focus on the ABCD dataset downloaded from NIH Data Archive (https://nda.nih.gov) [Casey et al., 2018]. The main goal of the ABCD study is to track the brain development from childhood through adolescence to understand biological and environmental factors that can affect the brain’s developmental trajectory. ABCD recruits approximately 10, 000 9-10 years old children. Longitudinal measures of the brain structure and function as well as rich behavior measures and genetic factors are collected across 21 sites in the United States [Auchter et al., 2018].

#### Imaging Data and Preprocessing

The ABCD imaging protocol is harmonized for three types of 3T scanners: Siemens Prisma, General Electric (GE) 750 and Philips. We downloaded the structural T1 MRI and diffusion MRI (dMRI) data for 5253 subjects from the ABCD 2.0 release. The structural T1 image was acquired with isotropic resolution of 1 *mm*^3^. The diffusion MRI image was obtained based on imaging parameters: 1.7 *mm*^3^ resolution, four different b-values (*b* = 500, 1000, 2000, 3000) and 96 diffusion directions. There are 6 directions at *b* = 500, 15 directions at *b* = 1000, 15 directions at *b* = 2000, 60 directions at *b* = 3000. Multiband factor 3 is used for dMRI acceleration. See Casey et al. [2018] for more details about the data acquisition and preprocessing of the ABCD data.

To obtain structural connectome, we used a state-of-the art dMRI data preprocessing framework named population-based connectome mapping (PSC) [Zhang et al., 2018]. PSC uses a reproducible probabilistic tractography algorithm [Girard et al., 2014; Maier-Hein et al., 2017] to generate the whole-brain tractography. This tractography method borrows anatomical information from high-resolution T1-weighted imaging to reduce bias in reconstruction of tractography. On average, about 10^6^ streamlines were generated for each subject. We used the popular Desikan-Killiany atlas [Desikan et al., 2006] to define the brain regions of interest (ROIs) corresponding to the nodes in the structural connectivity network. The Desikan-Killiany parcellation has 68 cortical surface regions with 34 nodes in each hemisphere. Freesurfer software [Dale et al., 1999; Fischl et al., 2004] is used to perform brain registration and parcellation. Next, for each pair of ROIs, we extracted the streamlines connecting them. In this process several procedures were used to increase the reproducibility: (1) each gray matter ROI is dilated to include a small portion of white matter region, (2) streamlines connecting multiple ROIs are cut into pieces so that we can extract the correct and complete pathway and (3) apparent outlier streamlines are removed. Extensive experiments have illustrated that these procedures can significantly improve the reproducibility of the extracted weighted networks, and readers can refer to Zhang et al. [2018] for more details. To analyze the brain as a network, a scalar number is usually extracted to summarize each connection. Here we use fiber count, but other measures, such as mean fractional anisotropy or connected surface area [Zhang et al., 2018] can be easily extracted using PSC.

Applying PSC, we processed 5253 subjects downloaded from NIH Data Archive. To control for handedness, we only focused on the 4213 right-handed subjects in our analysis. The distributions of the age, crystallized cognition composite score, picture vocabulary score and reading score are shown in Figure 2. For each composite or domain-specific crystallized intelligence score, subsets of subjects with age-adjusted scale scores 1 standard deviation above or below the national average are categorized into high vs. low crystallized intelligence group.

**Figure 2:**
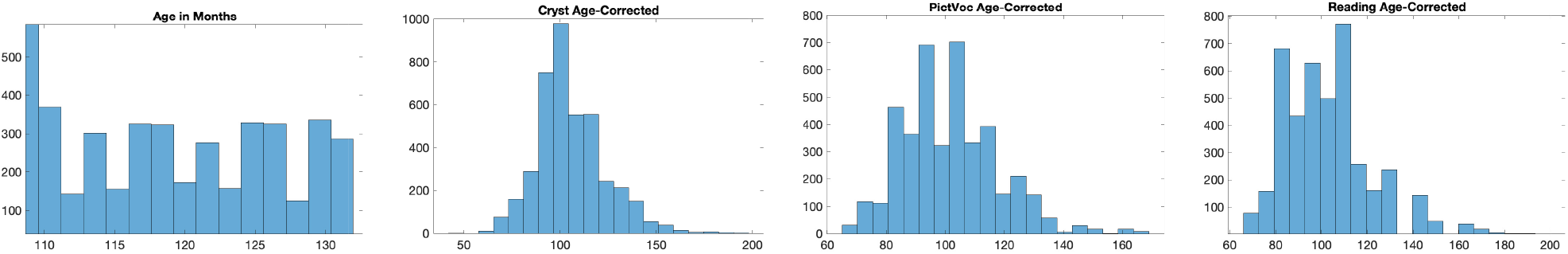
Histograms of ages, age-corrected crystallized cognition composite scores, picture vocabulary scores and reading scores of the 4213 subjects involved in this study.

### 2.2. Symmetric Bilinear Logistic Regression (SBLR)

Our data can be summarized as {(*y*_*i*_, ***x***_*i*_, *W*_*i*_) : *i* = 1, … , *n*}, where *y*_*i*_ is a binary response; ***x***_*i*_ ∈ ℝ^*m*^ contains the regular covariates of subject *i* (age, gender, etc.) with the first entry being 1; *W*_*i*_ denotes the structural brain network measured for subject *i*, which is a *V* × *V* symmetric matrix with zero diagonal entries. Our goal is to learn a set of small clique subgraphs from the brain network that are relevant to the outcome.

With this aim in mind, we propose the following symmetric bilinear logistic regression (SBLR):

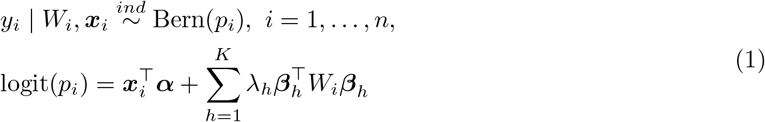

where ***α*** ∈ ℝ^*m*^ with the first entry being the intercept; ***β***_***h***_ ∈ ℝ^*V*^ and *λ*_*h*_ ∈ ℝ, *h* = 1, … , *K*. Model (1) assumes that the binary outcome *y*_*i*_ of each individual follows an independent Bernoulli distribution given the network observation *W*_*i*_ and other covariates ***x***_*i*_.

The coefficients of the network predictor in model (1) are assumed to have *K* components, where each component matrix 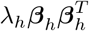 selects a signal subgraph. For ease of interpretation, the logit link of (1) can be written in the matrix dot product form:

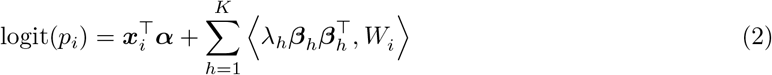

where ⟨*B*, *W*⟩ = trace(*B*^⊤^*W*) = vec(*B*)^⊤^vec(*W*). The parameters *λ*_*h*_’s in (2) are necessary to avoid constraining the coefficient matrix of *W*_*i*_ to be positive semi-definite. From (2), we can see that the nonzero entries in each component matrix 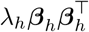 locate an outcome-relevant clique subgraph in the network predictor. If ***β***_*h*_ only contains two nonzero entries, the corresponding subgraph comes down to a single edge.

To ensure both identifiability of model (2) and sparsity of coefficient matrices 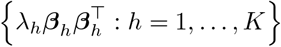, we penalize the magnitude of the lower-triangular entries in these coefficient matrices with the following elastic-net penalty:

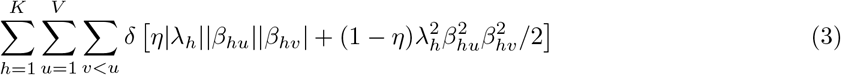

where the overall penalty factor *δ* > 0 and *η* ∈ [0, 1] controlling the fraction of *L*_1_ regularization.

### 2.3. Estimation

The parameters in SBLR model (1) are estimated by minimizing the loss function below:

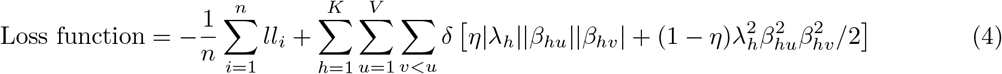

where *ll*_*i*_ is the log-likelihood of subject *i*. Plugging in the logit link of (1), we have

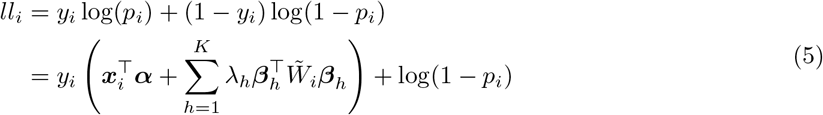

where 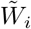 represents the normalized network observation with mean 0 and variance 1 for each edge.

The algorithm of block updating each component vector ***β***_*h*_ in tensor regression [Zhou et al., 2013] is not applicable to minimizing the loss function (4), because *ll*_*i*_ is not a concave function of ***β***_*h*_ when fixing the other parameters. Notice that

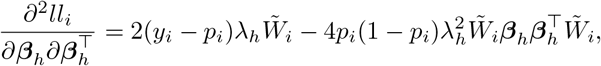

which may not be negative semi-definite. However, since the diagonal entries of each adjacency matrix 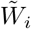 are zero, the loss function (4) is actually a convex function of each entry *β*_*hu*_ in ***β***_*h*_ when fixing the others, making the coordinate descent algorithm very appealing [Wang et al., 2019]. The challenge then lies in deriving the analytic form update for each parameter due to the nonsmoothness of (4). The technical details of coordinate descent algorithm are discussed below for model estimation.

#### 2.3.1. Updates for entries in 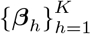

Minimizing the loss function (4) with respect to *β*_*hu*_, the *u*-th entry of ***β***_*h*_, given all the other parameters becomes:

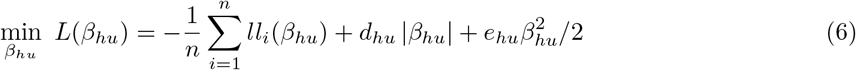

where

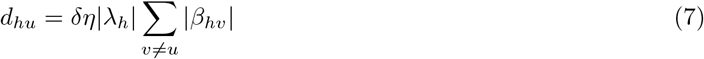

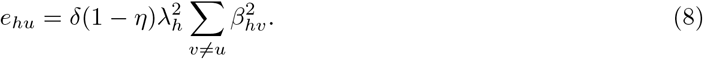

Equations (7) - (8) imply that the penalty factors *d*_*hu*_ and *e*_*hu*_ in (6) for |*β*_*hu*_| are related to the nonzero entries in ***β***_*h*_ excluding *β*_*hu*_. Hence *β*_*hu*_ is more likely to be shrunk to zero if the current number of nonzero entries in ***β***_*h*_ is large. This adaptive penalty will lead to a set of sparse vectors 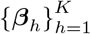 and hence a set of small signal subgraphs.

Note that

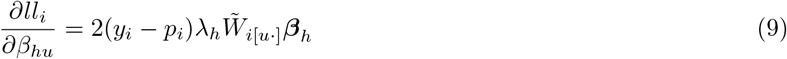

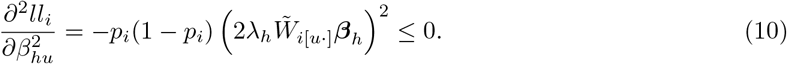

Therefore each log-likelihood *ll*_*i*_ is a concave function of *β*_*hu*_ when fixing the others, and (6) is a convex optimization for *β*_*hu*_.

Suppose the current estimate for *β*_*hu*_ at iteration *t* is 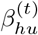 and fix the other parameters at their current estimates. The Newton algorithm for minimizing 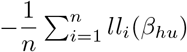 in (6) is equivalent to minimizing the following second-order Taylor expansion at the current estimate 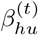:

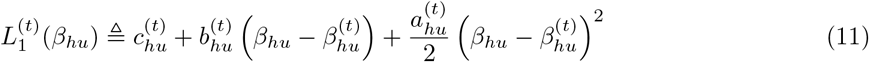

where

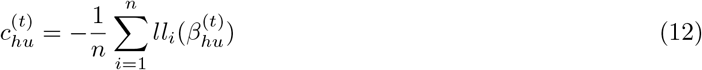

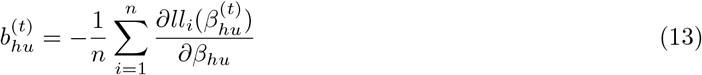

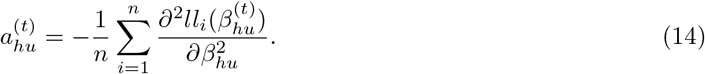

Similar to Friedman et al. [2010], *β*_*hu*_ is then updated by minimizing

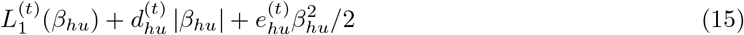

which has a closed form solution

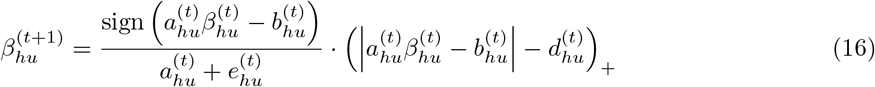

if 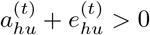. From (8), (10) and (14) we know that 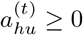 and 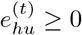.

The case 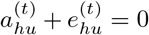 implies that (i) 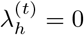 (ii) 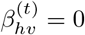 for *v* ≠ *u* or (iii) *η* = 1 and 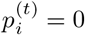 or 1, ∀*i*. For the former two cases, the lower triangular part of the component matrix 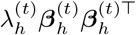 becomes zero no matter what value *β*_*hu*_ takes. So we set 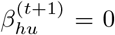 in these cases. Regarding (iii), if 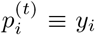, ∀*i*, (15) is minimized at *β*_*hu*_ = 0 because 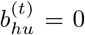 at this time according to (9) and (13) together with 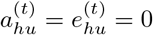. Then the first derivative of (15) becomes 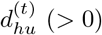 when *β*_*hu*_ > 0 and 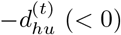 when *β*_*hu*_ < 0. Otherwise, 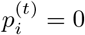 or 1 ∀*i* may be due to bad initialization. For example, a large magnitude of initial values of the parameters could easily make *p*_*i*_ become 1 or 0, ∀ i through the logit link in (1). In this case, setting *β*_*hu*_ = 0 could prevent the divergence of the solution. In practice, we always normalize the entries in {*W*_*i*_} before applying SBLR. We also recommend to initialize each parameter from *U* (−0.1, 0.1) to avoid the explosion in the logit scale.

The computational complexity of updating each entry *β*_*hu*_ is *O*(*nV*) per iteration and thus that of updating 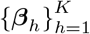 is *O*(*nKV*^2^).

#### 2.3.2. Updates for {λ_h_ : h = 1, … , K}

Minimizing the loss function (4) with respect to *λ*_*h*_ when fixing the others amounts to:

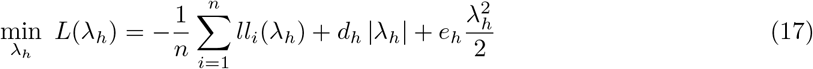

where 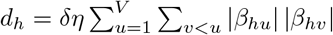 and 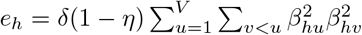.

Since *λ*_*h*_ is linear in the logit link (1) given the others, the log-likelihood is a concave function of *λ*_*h*_. Suppose the current estimate for *λ*_*h*_ is 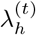. Similar to Section 2.3.1, *λ*_*h*_ is updated by minimizing the following quadratic approximation to (17):

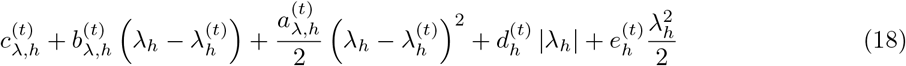

where

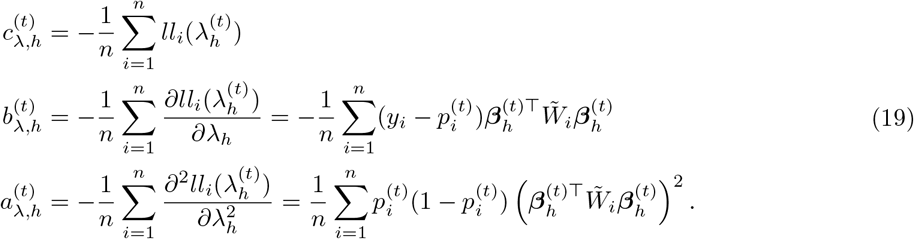

Note that 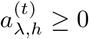 and 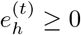. If 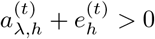, *λ*_*h*_ is updated to the argmin of (18):

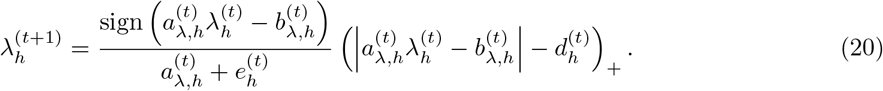

If 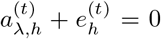, then either the component matrix 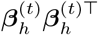 is a zero matrix or 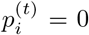 or 1, ∀*i*. In either case, 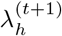 is set to 0 following a similar discussion in Section 2.3.1.

The computational complexity for updating each *λ*_*h*_ is *O*(*nV*^2^) per iteration and hence that of updating 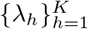 is *O*(*nKV*^2^).

#### 2.3.3. Update for α

***α*** ∈ ℝ^*m*^ is also updated by minimizing the quadratic approximation to 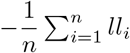 at the current estimate ***α***^(*t*)^ with the updating rule

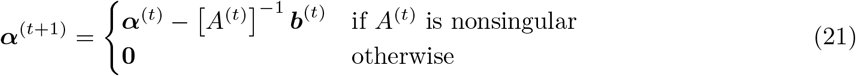

where

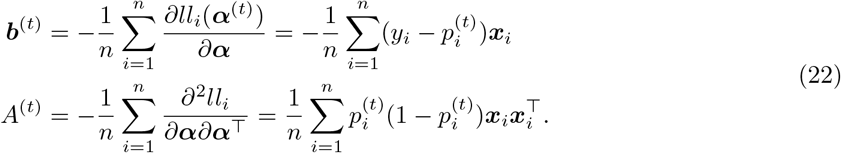

The computational complexity of this step is *O*(*m*^2^*n*).

**Algorithm 1.**
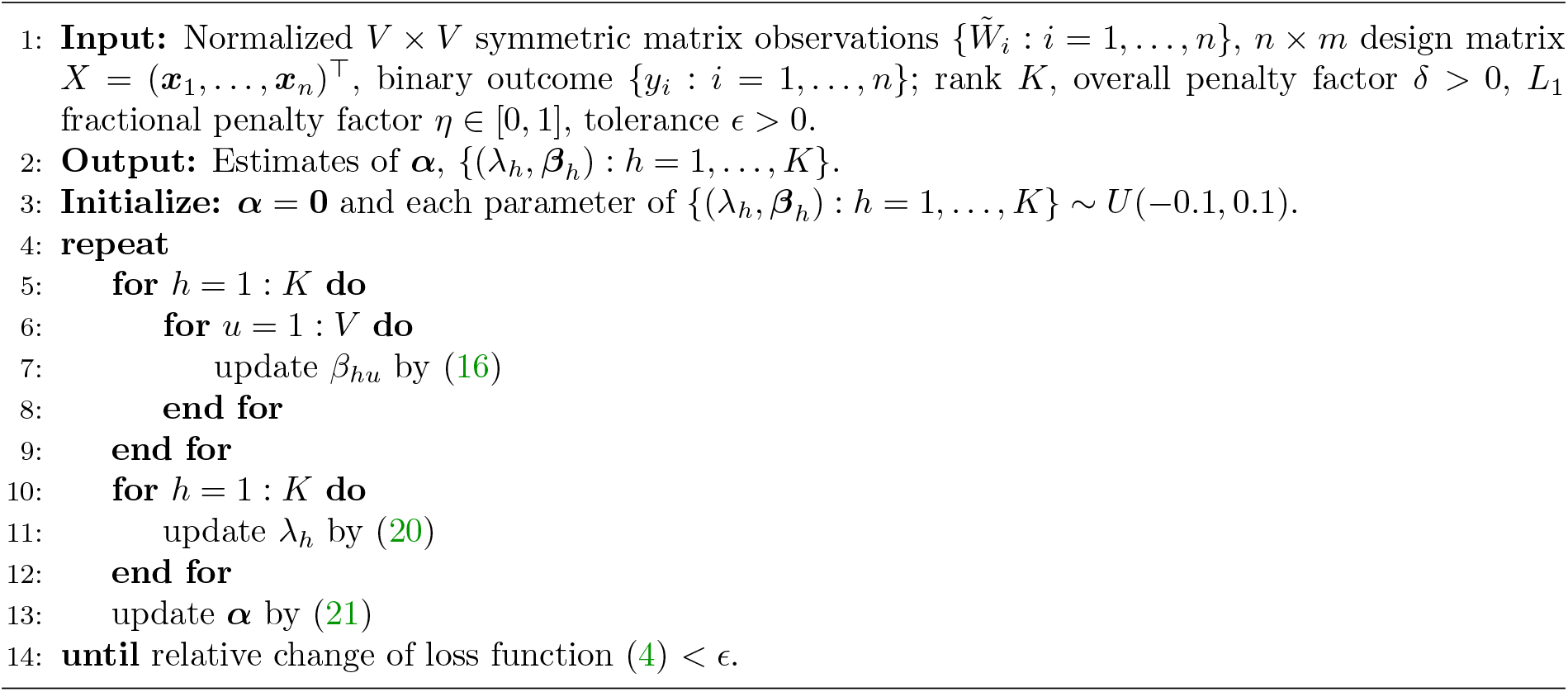
Coordinate descent for SBLR model with elastic-net penalty.

### 2.4. Other details

The above procedure is cycled through all the parameters until the relative change of the loss function (4) is smaller than a tolerance number *ϵ*, as summerized in Algorithm 1. Typical value is *ϵ* = 10^−5^ or 10^−6^. Since the loss function (4) is lower bounded by 0 and each update always decreases the function value, the coordinate descent algorithm derived above is guaranteed to converge.

In general, the algorithm should be run from multiple initializations to locate a good local solution. Although the estimates for (*λ*_*h*_, ***β***_*h*_) : *h* = 1, … , *K* will all be zero under sufficiently large penalty factor *δ*, we cannot initialize them at zero because the results will then get stuck at zero. The updating rules (16) and (20) imply that each parameter will be set to 0 given others being zero. In fact, we recommend to initialize all the parameters in (*λ*_*h*_, ***β***_*h*_) : *h* = 1, … , *K* to be nonzero in case some components get degenerated unexpectedly at the beginning. In practice, we initialize each parameter in (*λ*_*h*_, ***β***_*h*_) : *h* = 1, … , *K* from the uniform distribution *U* (−0.1, 0.1) as discussed at the end of Section 2.3.1.

### 2.5. Model selection

The penalty factor *δ* in the regularization (3) can be tuned on a validation set over a grid of values on [*δ*_min_, *δ*_max_] for a fixed *η*, where *δ*_max_ is a *roughly* smallest value for which all the parameters 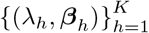 become zero, and *δ*_min_ = *εδ*_max_ (e.g. *ε* = 0.01). The optimal *δ* produces the smallest deviance (minus twice the log-likelihood) on validation set. The fractional parameter of *L*_1_ penalty, *η* ∈ [0, 1], can also be selected by validation.

Our proposed model (1) assumes a known number of components *K*. In practice, we choose a large enough number for *K* and allow the elastic-net penalty (3) to discard unnecessary components, leading to a data-driven estimate for the number of signal subgraphs. We assess the performance of the procedure and verify its lack of sensitivity to the chosen upper bound *K* in simulations.

## 3. Simulation Study

We use simulations to compare the performance of recovering true signal subgraphs and predictive performance among SBLR and the following methods:

1. Logistic regression with elastic-net penalty on vectorized networks, where the upper triangular entries of each adjacency matrix *W*_*i*_ are entered into a regularized logistic regression. The grid of values for the *L*_1_ fractional penalty factor *α* is picked as {0.1, 0.2, … , 1}. For each *α*, the optimal penalty factor *λ* is chosen from a sequence of 100 equally spaced *λ* values on the logarithmic scale. This method is fitted with `glmnet` toolbox in Matlab (http://www.stanford.edu/~hastie/glmnet_matlab).
2. Penalized graph classification (GC) approach [Relión et al., 2019], which also uses edge weights as predictors, but incorporates *L*_1_ and group lasso penalty to promote sparsity in selection of edges and nodes. Their penalty factor pair (*λ*, *ρ*) is tuned over a 5 × 11 grid, where *λ* ∈ {10^−4^, 10^−3^, … , 1} × *λ*_max_ and *ρ* ∈ {1, 10, 20, … , 100}, with (*λ*_max_, *ρ* = 100) ensuring that all the coefficients are penalized to zero. As Relión et al. [2019] suggest, values of *ρ* < 1 do not result in node selection. This method is fitted with graphclass package in R.
3. Screening method based on multiple testing with false discovery rate control (MT-FDR), where a two-sample *t*-test is performed on each edge in the network between the two groups. Multiple comparisons are adjusted by rejecting all local nulls having a *p*-value below the Benjamini-Yekutieli threshold [Benjamini et al., 2001] under arbitrary dependence assumptions on the multiple tests. The false discovery rate is controlled at level *α* = 0.05. A logistic regression is then fitted on the significant edge weights.
4. Tensor network factorization analysis (TNFA) [Zhang et al., 2019], which embeds the *V* × *V* symmetric adjacency matrices {*W*_*i*_ : *i* = 1, … , *n*} into a low dimensional *n* × *K* matrix *U* (*K* ≤ *V*). Each row *i* of *U* contains the *K* principal component scores of subject *i*, and each column of *U* corresponds to a basis network. A logistic regression is then fitted on the low embedding matrix *U*. The basis networks corresponding to the significant coefficients are selected as signal sub-networks. We use full rank *K* = *V* for TNFA in simulations, which explains around 99% variation in the generated networks on average.

In the data generating process, each adjacency matrix *W*_*i*_ is a 20 × 20 symmetric matrix generated from a set of 15 basis subgraphs with individual loading vectors

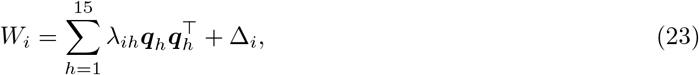

where ***q***_*h*_ ∈ {0, 1}^20^ is a random binary vector with ||***q***_*h*_||_0_ = *h* + 1, *h* = 1, 2, … , 15. The loadings {*λ*_ih_} in (23) are generated independently from *U* (0, 1). Δ_*i*_ is a symmetric 20 × 20 noise matrix that adds 5% noise to each edge in *W*_*i*_. The generating process (23) produces dense networks with complex correlation structure. These generated adjacency matrices *W*_*i*_ : *i* = 1, … , *n* are further standardized for each edge across subjects and the diagonals are set to zero.

The binary response *y*_*i*_ is associated with 3 subgraphs in the network. Specifically, *y*_*i*_ is generated from Bernoulli(*p*_*i*_) with

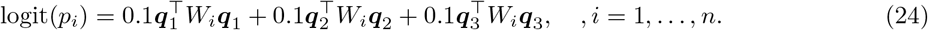

The generating process (24) indicates that the true signal subgraphs relevant to *y* correspond to the first three basis subgraphs 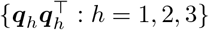 as displayed in Figure 3.

**Figure 3:**
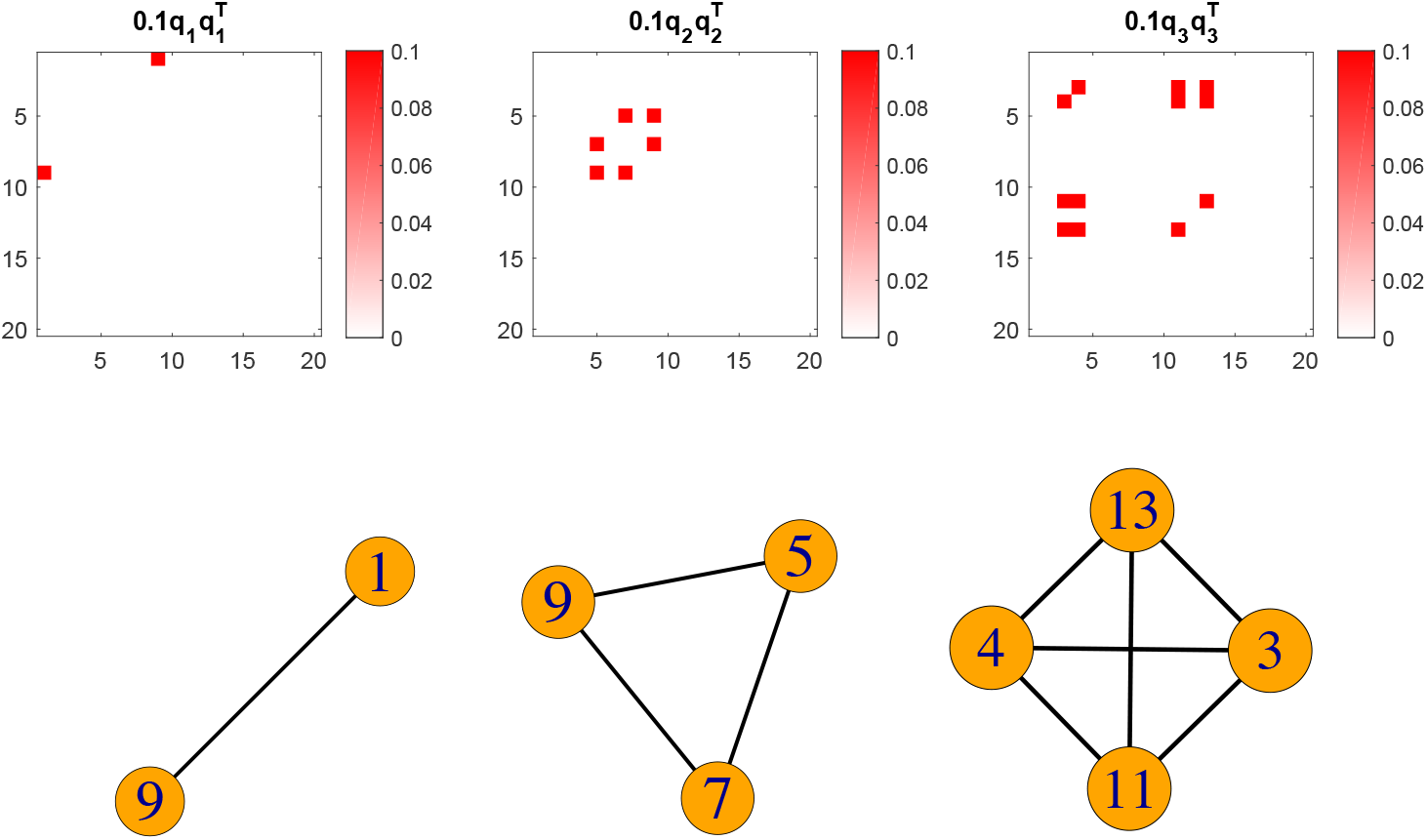
True signal subgraphs (lower panel) corresponding to 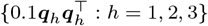 (upper panel) in one simulation.

We consider two sample sizes in this simulation: *n* = 500 and *n* = 1000. For ech sample size *n*, a dataset {(*W*_*i*_, *y*_*i*_) : *i* = 1, … , *n*} is drawn from the generating process (23) - (24). The dataset is then divided into two parts: training set (70%) and validation set (30%). Each method (SBLR, glmnet, GC) is fitted with training set and the optimal penalty pair is selected corresponding to the lowest deviance on validation set. We then refit the model with full dataset under the optimal penalty pair.

Some input parameters of Algorithm 1 for SBLR model are set as follows. The tolerance *ϵ* = 10^−6^ and *K* = 5. The *L*_1_ fractional penalty factor *η* is tuned over 5 fixed values {0.6, 0.7, … , 1}. For each value of *η*, we set *δ*_min_ = 0.01*δ*_max_ and choose a sequence of 11 equally spaced *δ* values on the logarithmic scale. 5 initializations are used in Algorithm 1.

The estimated results under optimal penalty pair from *glmnet* are displayed in Figure 4. As can be seen, the accuracy of glmnet improves as the sample size *n* increases, in terms of selecting more true signal edges and fewer non-signal edges. But it is difficult to identify meaningful structure among the selected edges. Figure 5 displays the estimated results under optimal penalty pair for the *GC* approach [Relión et al., 2019]. Although this approach identifies all the true signal edges under larger sample size, it also selects a larger number of false edges. As with glmnet, this approach does not guarantee any structure among the selected edges.

**Figure 4:**
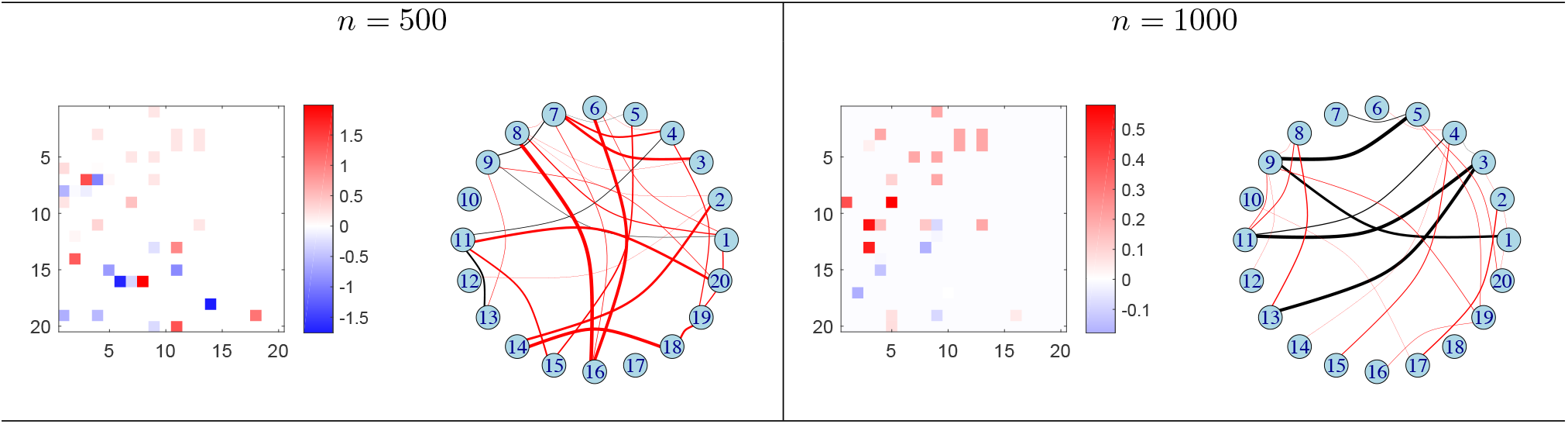
Estimated results of glmnet under *n* = 500 and *n* = 1000. In each panel, the left plot shows estimated coefficients (lower triangular) versus true values (upper triangular); the right plot shows the selected edges in the network, where black edges denote true signal edges and red ones falsely identified edges; the thickness of each edge is proportional to the magnitude of its estimated coefficient.

**Figure 5:**
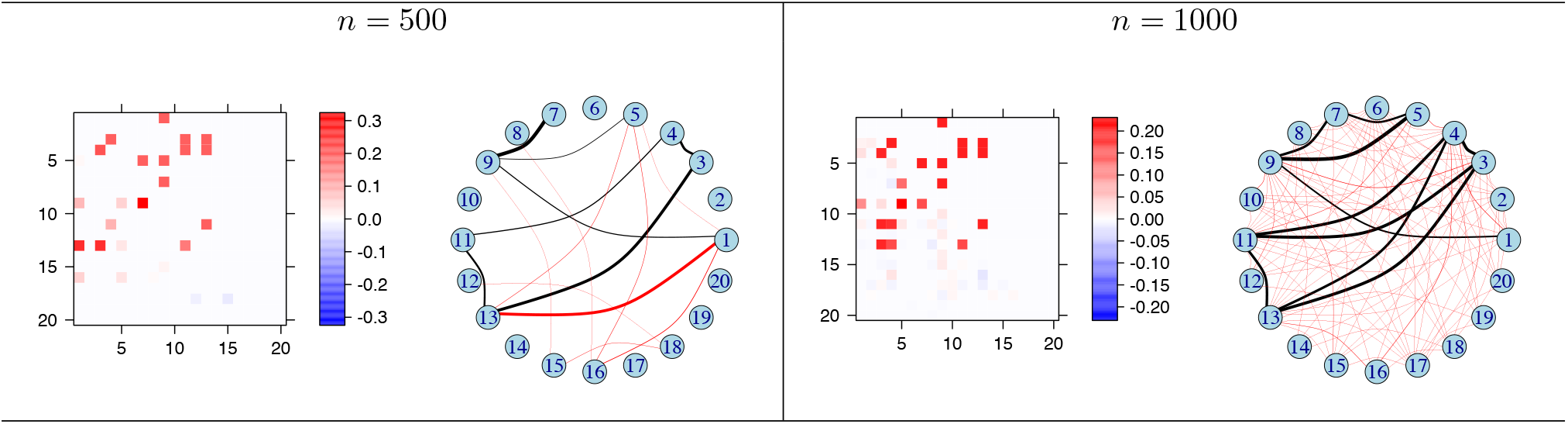
Estimated results of penalized graph classification (GC) approach under *n* = 500 and *n* = 1000. In each panel, the left plot shows estimated coefficients (lower triangular) versus true values (upper triangular); the right plot shows the selected edges in the network, where black edges denote true signal edges and red ones falsely identified edges; the thickness of each edge is proportional to the magnitude of its estimated coefficient.

Multiple *t*-test screening method (MT-FDR) deems many more edges to be significant in this case, taking up about 86% and 95% of the total number of edges in the network under *n* = 500 and *n* = 1000 respectively. Tensor factorization approach (TNFA) on the contrary selects no basis networks under each sample size, because none of the 20 components are significant in the logistic regression at the significance level of 0.05.

The estimated results under optimal penalty pair of our *SBLR* under *n* = 500 and *n* = 1000 are displayed in Figure 6. The performance of SBLR improves with larger sample size. Compared to Figure 3, Figure 6 shows that SBLR recovers all the true signal subgraphs under *n* = 1000 with fewer wrong edges compared to *n* = 500.

**Figure 6:**
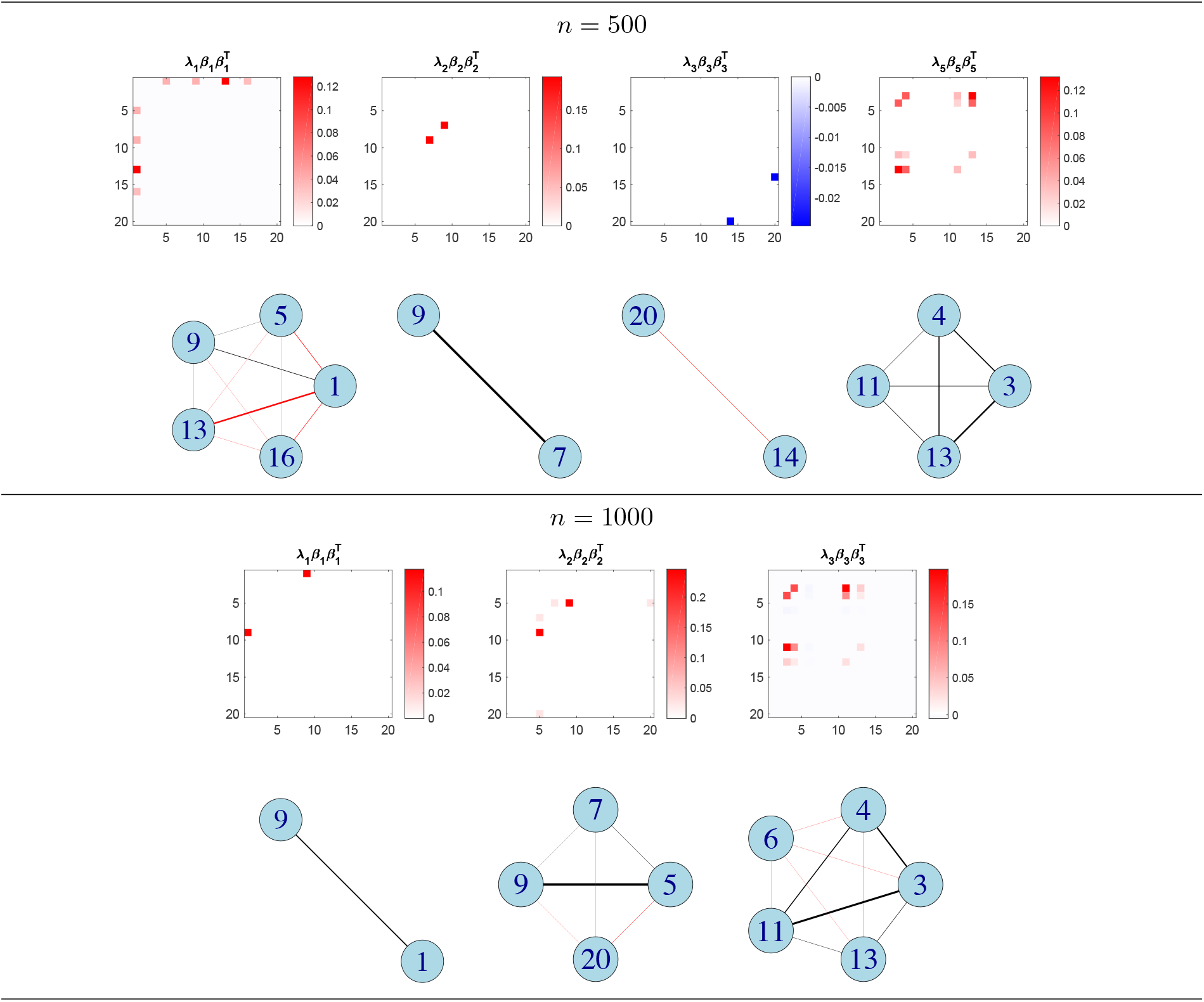
Estimated nonzero coefficient components 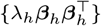 from SBLR (upper) and their selected subgraphs (lower) under *n* = 500 and *n* = 1000. Black edges denote true signal edges and red ones falsely identified edges; the thickness of each edge is proportional to the magnitude of its estimated coefficient.

The procedure above is repeated 100 times. For either sample size *n*, 100 datasets are generated based on the generating process (23) - (24). For each dataset, we additionally generate observation pairs {(*W*_*i*_, *y*_*i*_)} for 200 subjects as test data and compute the area under an ROC curve (AUC) [DeLong et al., 1988] and deviance [McCullagh and Nelder, 1989] on test data for each method. In addition, we record for each method the true positive rate (TPR) representing the proportion of true signal edges that are correctly identified, and the false positive rate (FPR) representing the proportion of non-signal edges that are falsely identified. Performances under two different choices of *K* are compared for SBLR: *K* = 5 and *K* = 10. Table 1 displays the mean and standard deviation (SD) of the TPR, FPR, AUC and deviance on test data for each method.

**Table 1:** Mean and standard deviation of TPR, FPR, AUC and deviance on test data across 100 simulations. *Glmlasso* is logistic regression with *L*_1_ penalty; SBLR with *η* = 1 also indicates that only *L*_1_ penalty is applied.

|  | TPR | FPR | AUC | deviance |
| --- | --- | --- | --- | --- |
| $n = 500$ | | | | |
| glmlasso | 0.5766±0.1423 | **0.0516±0.0693** | 0.7931±0.0366 | 1.1061±0.0783 |
| glmnet | 0.7034±0.1987 | 0.1208±0.1688 | 0.7931±0.0359 | 1.1061±0.0762 |
| GC | 0.8327±0.1585 | 0.2286±0.2649 | 0.7942±0.0355 | **1.1026±0.0752** |
| MT-FDR | **0.9940±0.0278** | 0.8953±0.0706 | 0.6752±0.0408 | 3.6586±1.7923 |
| TNFA | 0.1000±0.3015 | 0.1000±0.3015 | 0.7865±0.0352 | 1.1252±0.0856 |
| SBLR ($K = 5, \eta = 1$) | 0.7139±0.1770 | 0.0738±0.0601 | 0.7941±0.0347 | 1.1032±0.0718 |
| SBLR ($K = 5$) | 0.7451±0.1783 | 0.0835±0.0621 | 0.7940±0.0355 | 1.1031±0.0736 |
| SBLR ($K = 10, \eta = 1$) | 0.7142±0.1662 | 0.0709±0.0450 | **0.7946±0.0358** | **1.1026±0.0739** |
| SBLR ($K = 10$) | 0.7442±0.1904 | 0.0921±0.0729 | 0.7937±0.0355 | 1.1041±0.0746 |
| $n = 1000$ | | | | |
| glmlasso | 0.6830±0.1463 | **0.0502±0.0525** | 0.8032±0.0336 | 1.0823±0.0726 |
| glmnet | 0.8062±0.1804 | 0.1279±0.1641 | 0.8035±0.0337 | 1.0818±0.0726 |
| GC | 0.9204±0.1240 | 0.3172±0.3104 | 0.8035±0.0334 | 1.0818±0.0731 |
| MT-FDR | **0.9990±0.0100** | 0.9478±0.0366 | 0.7437±0.0403 | 1.3821±0.1461 |
| TNFA | 0.1800±0.3861 | 0.1800±0.3861 | 0.7985±0.0322 | 1.0906±0.0769 |
| SBLR ($K = 5, \eta = 1$) | 0.8221±0.1580 | 0.0717±0.0476 | 0.8037±0.0332 | 1.0815±0.0709 |
| SBLR ($K = 5$) | 0.8704±0.1377 | 0.0812±0.0516 | 0.8039±0.0333 | 1.0813±0.0711 |
| SBLR ($K = 10, \eta = 1$) | 0.8101±0.1529 | 0.0739±0.0429 | 0.8037±0.0329 | 1.0810±0.0710 |
| SBLR ($K = 10$) | 0.8510±0.1298 | 0.0810±0.0512 | **0.8041±0.0335** | **1.0806±0.0718** |

Table 1 shows that the mean and SD of the four measures are very similar for SBLR under *K* = 5 and *K* = 10, with the same type of penalty (*L*_1_ or elastic-net) and the same sample size. This implies that the performance of SBLR is robust to the chosen upper bound for the rank. In addition, Table 1 shows that the mean TPR, AUC and deviance improve with larger sample size for all the methods, but the mean FPRs of GC, multiple testing (MT-FDR) and tensor factorization (TNFA) increase considerably when *n* goes from 500 to 1000. The mean FPRs of GC and MT-FDR are also much higher than that of the other methods. All the methods except for MT-FDR have competitive predictive performance in terms of similar AUC and deviance on test data. Glmlasso has the lowest FPR on average but also low mean TPR. SBLR models with either *L*_1_ or elastic-net penalty achieve higher TPR and lower FPR on average than glmnet does under either sample size.

All the numerical experiments are conducted on a machine with six Intel Core i7 3.2 GHz processors and 64 GB of RAM. Algorithm 1 of SBLR method is implemented in Matlab (R2018a). Under the sample size *n* = 1000, SBLR with *K* = 10 takes 99.6 seconds on average to run a validation instance over the 5 × 11 grid, glmnet takes 21 seconds and GC approach takes 130.4 seconds on average.

## 4. Application: SC Subgraphs Distinguishing High and Low Crystallized Intelligence of Adolescents

We apply our method to the ABCD dataset described in Section 2.1 to examine the associations between structural brain network and crystallized intelligence for adolecents aged 9-10 years. NIH toolbox measures crystallized intelligence through two core tests: Picture Vocabulary test and Oral Reading Recognition test. They also provide a composite score, Crystallized Cognition Composite, to allow for evaluation of overall crystallized intelligence.

SBLR model is applied to differentiate high from low crystallized intelligence for each composite or domain-specific score, as well as identifying clique signal subgraphs contributing to the classification. Some input parameters of Algorithm 1 for SBLR are set below in this section. The *L*_1_ fractional penalty factor *η* is set to 1 to encourage sparsity in the results, as the AUC on test data is not sensitive to different values of *η* in this case. The tolerance *ϵ* = 10^−5^ and *K* = 20. 5 initializations are used in Algorithm 1. SBLR is trained over a sequence of 11 equally spaced *δ* values on the logarithmic scale with *δ*_min_ = 0.1*δ*_max_. We also compare to glmlasso, multiple testing screening (MT-FDR) and tensor factorization analysis (TNFA) on variable selection and predictive performance. Full rank *K* = 68 is set in TNFA, which explains around 75% of the variation in the brain networks.

We employ stability selection [Meinshausen and Bühlmann, 2010; Shah and Samworth, 2013] to enhance significance of the selected variables (subgraphs or individual edges) for each method, which involves many rounds of random data splitting and keeping the variables with high selection probability. Specifically, the dataset for each cognitive measure is divided into 3 parts: training (60%), validation (20%) and test set (20%). SBLR and glmlasso are fitted with training set and the optimal *L*_1_ penalty factor is tuned by validation set. MT-FDR and TNFA are fitted with training and validation sets. We record the selected variables for each method (under the optimal penalty factor) as well as AUC on test data across 30 rounds of random data splitting, and calculate the probability of each variable being selected.

### 4.1. Picture Vocabulary

The Picture Vocabulary test uses an audio recording of words, presented with four photographic images on the computer screen. The participants are asked to select the picture that best matches the meaning of the word. Using the 1 standard deviation rule proposed in section 2.1, we obtain a dataset containing 1034 kids with high picture vocabulary scores and 282 low scores.

SBLR on average selects 9.6 nonzero subgraphs 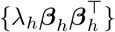 across 30 data splittings with a standard deviation of 4.0. Summarizing the results from stability selection [Meinshausen and Bühlmann, 2010; Shah and Samworth, 2013], the corresponding subgraphs with more than two ROIs and selection probabilities greater than 0.5 are displayed in Figure 7. Figure 7 also displays individual connections selected by glmlasso and MT-FDR with probabilities greater than 0.5. Note that the coefficients with large magnitude in the subgraphs identified by SBLR look similar to the estimates from glmlasso and MT-FDR. But the connections selected by the latter two methods lack meaningful structure and are difficult to justify neurologically. 12 components remain significant in TNFA more than 50% of the time, but their corresponding basis networks involve all the connections in the brain network. The average AUCs on test data across 30 data splittings are all around 0.78 for SBLR, glmlasso and TNFA with a standard deviation around 0.03, while that of MT-FDR is 0.72 ± 0.04.

**Figure 7:**
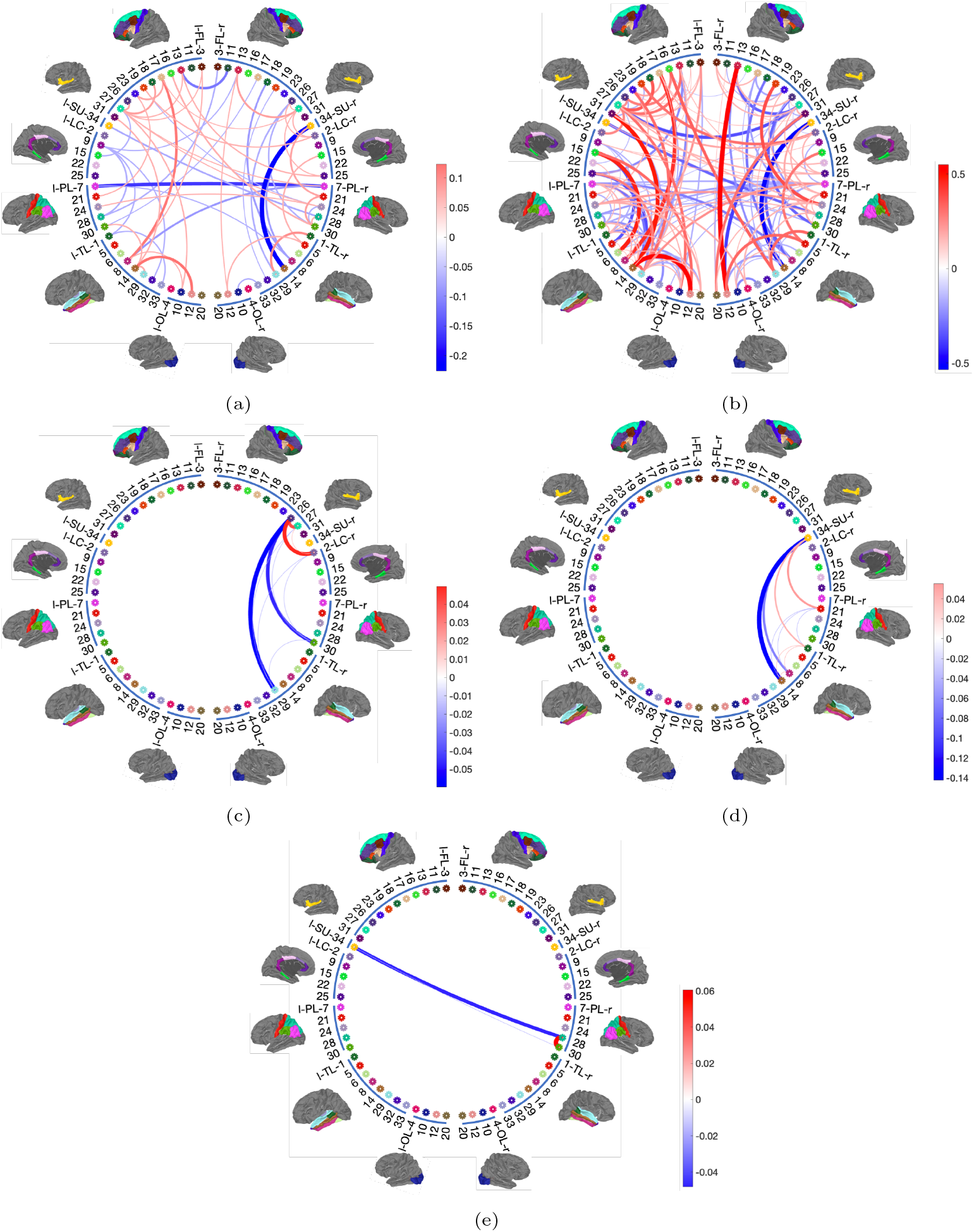
Results of Picture Vocabulary: (a) connections selected by glmnet with selection probabilities > 0.5 across 30 rounds of random data splitting; (b) connections selected by MT-FDR with selection probabilities > 0.5; (c) - (e) subgraphs (≥ 3 ROIs) selected by SBLR with selection probabilities > 0.5. The subgraphs are arranged in the descending order of selection frequency. The thickness of each edge is proportional to the magnitude of its mean estimated coefficient; the color goes from blue to red as the coefficient goes from negative to positive.

The three subgraphs identified in Figure 7 all seem to have hub structure (if we only consider large connections), with the hub nodes being 26*r* (right rostral middle frontal), 34*r* (right insula) and 28*r* (right superior parietal) respectively. Plot (c) of Figure 7 implies that right-handed children with stronger neural connections among 26*r* (right rostral middle frontal), 27*r* (right superior frontal) and 2*r* (right caudal anterior cingulate), and weaker connections among 26*r*, 29*r* (right superior temporal) and 30*r* (right supramarginal) are more likely to get high scores on Picture Vocabulary test. Plot (d) of Figure 7 implies that right-handed children with stronger neural connections among 34*r* (right insula), 21*r* (right postcentral) and 1*r* (right bankssts), and weaker connections among 34*r*, 14*r* (right middle temporal) and 8*r* (right inferior temporal) are more likely to get high scores in Picture Vocabulary test. Plot (e) of Figure 7 implies that right-handed children with stronger neural connection between 28*r* (right superior parietal) and 30*r* (right supramarginal), and weaker connection between 28*r* (right superior parietal) and 34*l* (left insula) are more likely to achieve high ability of this cognition measure. Some of these brain regions agree with the findings in neuroscience literature, for example, right superior parietal gyrus, right supramarginal gyrus and left insula are among the activated regions in auditory language processing tasks for children [Vogan et al., 2016; Sugiura et al., 2011; Oh et al., 2014].

### 4.2. Reading Recognition

Participants on this test are asked to read and pronounce letters and words as accurately as possible. Applying the 1 standard deviation rule proposed in section 2.1, we construct a dataset containing 918 subjects with high Reading Recognition scores and 477 subjects with low scores.

SBLR on average selects 9.3 ± 4.2 nonzero subgraphs across 30 data splittings. Figure 8 displays the subgraphs identified by SBLR with selection probabilities > 0.5, along with connections selected by glmlasso and MT-FDR with probabilities > 0.5. Most estimated coefficients with large magnitude in SBLR selected subgraphs look similar to those from glmlasso and MT-FDR. Four components remain significant in TNFA more than 50% times, which correspond to all the connections in the brain network. The average AUCs on test data across 30 splittings are approximately 0.7 for all the methods above with a standard deviation around 0.03.

**Figure 8:**
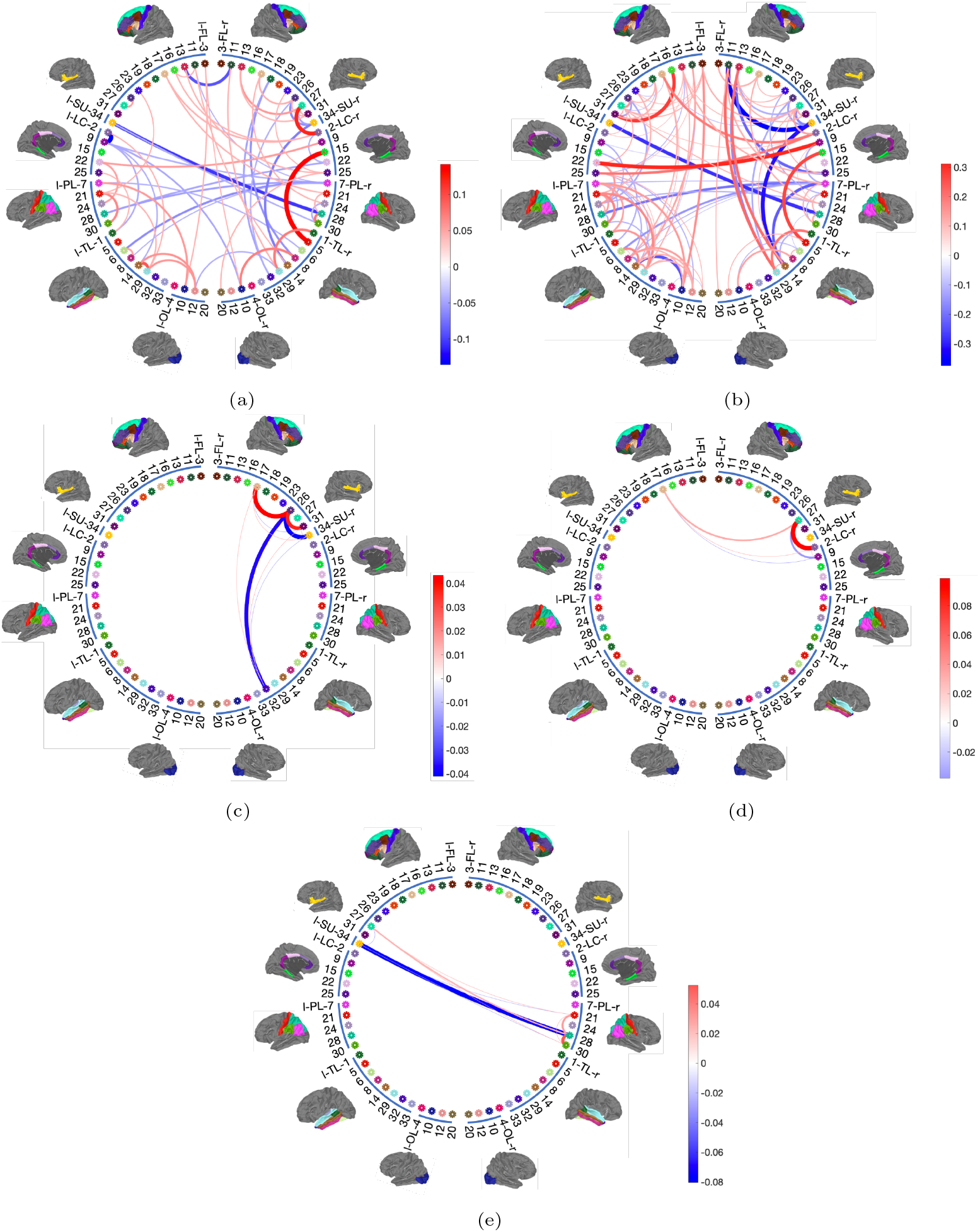
Results of Reading Recognition: (a) connections selected by glmnet with selection probabilities > 0.5 across 30 dataset splittings; (b) connections selected by MT-FDR with selection probabilities > 0.5; (c) - (e) subgraphs (≥ 3 ROIs) selected by SBLR with selection probabilities > 0.5. The subgraphs are arranged in the descending order of selection frequency. The thickness of each edge is proportional to the magnitude of its mean estimated coefficient; the color goes from blue to red as the coefficient goes from negative to positive.

The subgraphs in Figure 8 located by SBLR display hub structure again and share many similarities to those associated with Picture Vocabulary. Compared to Figure 7, Reading Recognition measure has two hub nodes in common with Picture Vocabulary: 26*r* (right rostral middle frontal) and 28*r* (right superior parietal). And Plot (e) of Figure 7 is a subgraph of Plot (e) in Figure 8. The latter has two extra ROIs: 21*r* (right postcentral) and 27*l* (left superior frontal), and stronger connections among the two nodes and the hub node (28*r*) have positive effect on the cognitive functioning of Reading Recognition. Plot (c) of Figure 8 implies that right-handed children with stronger neural connections among 17*r* (right pars opercularis), 26*r* (right rostral middle frontal) and 31*r* (right frontal pole) are more likely to get high scores on Reading Recognition test, while the connection strengths among 26*r*, 32*r* (right temporal pole) and 34*r* (right insula) may have negative effect on this cognitive functioning. Plot (d) of Figure 8 implies that stronger neural connections among 17*l* (left pars opercularis), 27*r* (right superior frontal) and 2*r* (right caudal anterior cingulate) have positive effect on this cognitive functioning, while the connection strength between 27*r* and 9*r* (right isthmus cingulate) may have a slight negative effect.

### 4.3. Crystallized Cognition Composite

Crystallized Cognition Composite can be interpreted as a global assessment of verbal reasoning. This composite score is derived by averaging the normalized scores of Picture Vocabulary and Reading Recognition test, and the age-adjusted scale scores are calculated based on this new distribution [Weintraub et al., 2013]. Under the 1 standard deviation rule, we identify 1084 high crystallized cognitive cases and 536 low crystallized cognitive cases.

SBLR on average selects 12.3 ± 3.5 nonzero subgraphs across 30 data splittings. Figure 9 displays the subgraphs identified by SBLR with selection probabilities > 0.5, along with individual connections selected by glmlasso and MT-FDR with probabilities > 0.5. 9 components maintain significant in TNFA more than 50% times, which again corresponds to all the connections in the brain network. The AUCs on test data across 30 data splittings are around 0.76 ± 0.02 for SBLR, glmlasso and TNFA, while that of MT-FDR is 0.71 ± 0.03.

**Figure 9:**
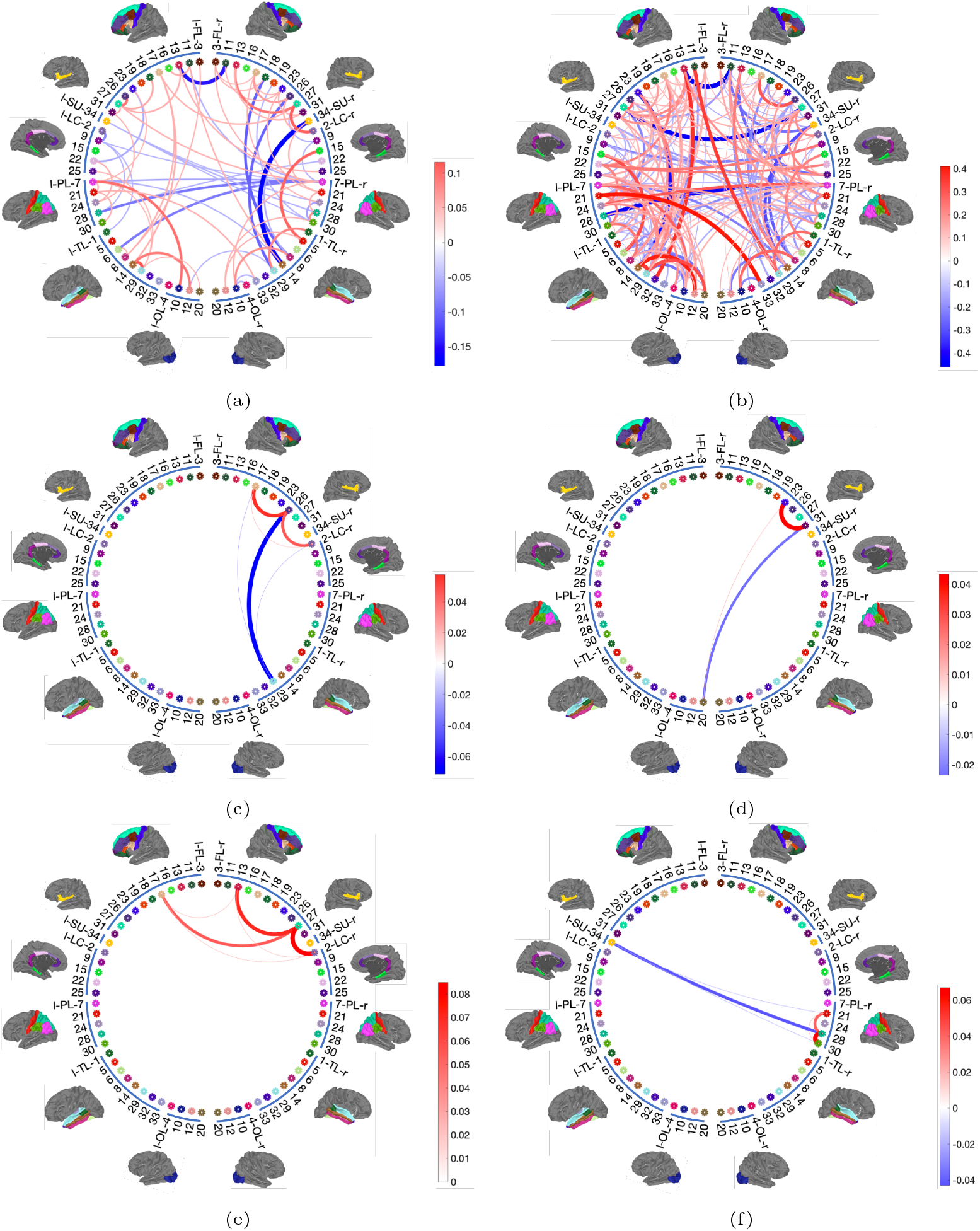
Results of Crystallized Cognition Composite: (a) connections selected by glmnet with selection probabilities > 0.5 across 30 dataset splittings; (b) connections selected by MT-FDR with probabilities > 0.5; (c) - (f) subgraphs (≥ 3 ROIs) selected by SBLR with probabilities > 0.5. The subgraphs are arranged in the descending order of selection frequency. The thickness of each edge is proportional to the magnitude of its mean estimated coefficient; the color goes from blue to red as the coefficient goes from negative to positive.

The subgraphs identified by SBLR in Figure 9 predictive of high and low crystallized cognitive ability have many overlaps with the selected subgraphs associated with Picture Vocabulary and Reading Recognition. The subgraph in Plot (c) of Figure 9 seems like a “composite” of the subgraphs in Plot (c)’s of both Figure 7 and Figure 8. Plot (f) of Figure 9 echoes Plot (e)’s in both Figure 7 and Figure 8. The subgraph in Plot (e) of Figure 9 overlaps with that in Plot (d) of Figure 8. Plot (e) of Figure 9 implies that right-handed children with stronger neural connections among 17*l* (left pars opercularis), 13*r* (right medial orbitofrontal), 27*r* (right superior frontal) and 2*r* (right caudal anterior cingulate) are more likely to have high level of functioning on crystallized cognition. Plot (d) of Figure 9 implies that stronger connection between 31*r* (right frontal pole) and 23*r* (right precentral) has positive effect on crystallized cognitive functioning, while the connection strength between 31*r* and 20*l* (left pericalcarine) may have negative effect.

## 5. Discussion

We have presented a useful tool for studying differences in brain connectivity patterns between groups, which produces more interpretable results than unstructured classifiers do, while maintaining competitive predictive performance. Simulation studies show that the selection accuracy of SBLR could generally be improved by increasing the number of subjects in the data.

Each subgraph identified by SBLR is a clique, which is a fully connected subgraph among a subset of nodes. While such a clique subgraph can be biologically meaningful, it is also restrictive - the number of connections in the subgraph increases at the order of *O*(*m*^2^) for *m* selected nodes. However, SBLR generally does not select very large subgraphs under the elastic-net penalty. There are other ways to define subgraphs, for example, Vogelstein et al. [2012] defines a subgraph as a minimum set of vertices and edges distinguishing groups, and Khambhati et al. [2018] constructed subgraphs based an unsupervised non-negative matrix factorization, which is similar to Zhang et al. [2019]. More flexible ways of defining subgraphs with more interesting network topological properties can be explored in the future.

Very few studies have focused on WM’s contribution to cognition. Specific to crystallized cognition, there have only been a handful of studies examining major WM tracts’ roles. For example, higher FA in superior longitudinal fasciculus was related to higher crystallized cognition in children [Simpson-Kent et al., 2020], higher FA in forceps minor was related to higher crystallized cognition in adults [Góngora et al., 2020]. Both tracts contain numerous inter-hemisphere connections across several brain regions (e.g., superior longitudinal fasciculus) or within frontal region (e.g., forceps minor). Our study represents the very first effort in the literature specifying clique subgraphs of SC related to crystallized cognition. Importantly, across composite score and individual domains of crystallized cognition, we identified consistent brain regions and subgraphs (refer to Figure 7, 8 and 9) predominantly in right-hemisphere frontal-parietal regions. The finding is consistent to cumulative functional subgraph literature on frontal-parietal driven executive network during neuro-development [Chai et al., 2017], and flexible periphery of the language network [Fedorenko and Thompson-Schill, 2014].

We also noticed that the classification accuracy with AUC around 0.78 for crystallized cognition is very high. From our previous study, SC is robust and reproducible and more predictive of cognition compared with functional connectivity (FC) derived from functional MRI [Zhang et al., 2018, 2019]. We hypothesize that SC is a better biomarker for understanding the cognition development in adolescents. To verify this hypothesis, analyses and comparisons with FC seem to be a natural next step.

The frontal-parietal driven subgraphs among those aged 9-10 years represent a set of critical biomarkers for overall neuro-development. However, these subgraphs’ physiological meaning can be transient across ages. Comparisons across a broader age range should be conducted to further confirm the role of these subgraphs in crystallized cognition and its development. With more data being recorded in the ABCD study, we hope to further analyze SC and cognition development.

## 6. Acknowledgement

Data used in the preparation of this article were obtained from the Adolescent Brain Cognitive Development (ABCD) Study (https://abcdstudy.org), held in the NIMH Data Archive (NDA). This is a multisite, longitudinal study designed to recruit more than 10,000 children age 9-10 and follow them over 10 years into early adulthood. The ABCD Study is supported by the National Institutes of Health and additional federal partners under award numbers U01DA041022, U01DA041028, U01DA041048, U01DA041089, U01DA041106, U01DA041117, U01DA041120, U01DA041134, U01DA041148, U01DA041156, U01DA041174, U24DA041123, U24DA041147, U01DA041093, and U01DA041025. A full list of supporters is available at https://abcdstudy.org/federal-partners.html. A listing of participating sites and a complete listing of the study investigators can be found at https://abcdstudy.org/scientists/workgroups/. ABCD consortium investigators designed and implemented the study and/or provided data but did not necessarily participate in analysis or writing of this report. This manuscript reflects the views of the authors and may not reflect the opinions or views of the NIH or ABCD consortium investigators.

The research of L. Wang is supported by National Natural Science Foundation of China [grant number 11901583]. Zhang wants to acknowledge support from the National Institutes of Health (NIH) of the United States under award number MH118927 and MH118020.

